# Impact of Screening and Follow-up Colonoscopy Adenoma Sensitivity on Colorectal Cancer Screening Outcomes in the CRC-AIM Microsimulation Model

**DOI:** 10.1101/2020.08.31.271924

**Authors:** Deborah A. Fisher, Leila Saoud, Kristen Hassmiller Lich, A. Mark Fendrick, A. Burak Ozbay, Bijan J. Borah, Michael Matney, Marcus Parton, Paul J. Limburg

## Abstract

**Background:** Real-world data for patients with positive colorectal cancer (CRC) screening stool-tests demonstrates that adenoma detection rates are lower when endoscopists are blinded to the stool-test results. This suggests adenoma sensitivity may be lower for screening colonoscopy than for follow-up to a known positive stool-based test. Previous CRC microsimulation models assume identical sensitivities between screening and follow-up colonoscopies after positive stool-tests. The Colorectal Cancer and Adenoma Incidence and Mortality Microsimulation Model (CRC-AIM) was used to explore the impact on screening outcomes when assuming different adenoma sensitivity between screening and combined follow-up/surveillance colonoscopies.

**Methods:** Modeled screening strategies included colonoscopy every 10 years, triennial multitarget stool DNA (mt-sDNA), or annual fecal immunochemical test (FIT) from 50-75 years. Outcomes were reported per 1,000 individuals without diagnosed CRC at age 40. Base-case adenoma sensitivity values were identical for screening and follow-up/surveillance colonoscopies. Ranges of adenoma sensitivity values for colonoscopy performance were developed using different slopes of odds ratio adjustments and were designated as small, medium, or large impact scenarios.

**Results:** As the differences in adenoma sensitivity for screening versus follow-up/surveillance colonoscopies became greater, life-years gained (LYG) and reductions in CRC-related incidence and mortality versus no screening increased for mt-sDNA and FIT and decreased for screening colonoscopy. The LYG relative to screening colonoscopy reached >90% with FIT in the base-case scenario and with mt-sDNA in a “medium impact” scenario.

**Conclusions:** Assuming identical adenoma sensitivities for screening and follow-up/surveillance colonoscopies underestimates the potential benefits of stool-based screening strategies.

## Introduction

Screening average-risk individuals for colorectal cancer (CRC) is recommended by the American Cancer Society (ACS), ^1^ US Preventive Services Task Force (USPSTF),^2^ and other national organizations. Screening modalities include colonoscopy and stool-based tests, such as fecal immunochemical testing (FIT) and the multi-target stool DNA (mt-sDNA) assay, among other endorsed options. The results of CRC screening microsimulation models developed by the Cancer Intervention and Surveillance Modeling Network (CISNET) Colorectal Working Group have helped inform screening guidelines by modeling outcomes from various screening modalities over a range of patient ages and testing intervals.^3–8^

For simplicity, CRC microsimulation models generally assume that the sensitivity of follow-up colonoscopy for adenoma detection after a positive stool-based test is the same as with screening colonoscopy.^3,4^ However, this simplistic assumption does not align with real-world observations. Several previous studies have demonstrated that the adenoma detection rate (ADR; proportion of patients with ≥1 detected adenoma) is approximately 20% higher when a follow-up colonoscopy is performed for the indication of a positive stool-based test compared with a primary screening colonoscopy exam.^9–12^ The higher yield for adenomas at follow-up colonoscopy may be related to patient biology, endoscopist performance, or a combination of both. A previous study showed that endoscopists who were aware of a positive mt-sDNA screening result (unblinded) had a longer colonoscopy withdrawal time, found significantly more adenomas, and had a higher ADR compared with those endoscopists who were not aware of a positive test (blinded).^13^ To more accurately model the differential clinical outcomes between screening and follow-up colonoscopy exams, we used the validated Colorectal Cancer and Adenoma Incidence and Mortality Microsimulation Model (CRC-AIM)^14^ to simulate the impact on estimated outcomes, including CRC incidence, CRC mortality, and life-years gained (LYG), as well as adenoma miss rate (AMR) and ADR when assuming different adenoma sensitivity between screening and combined follow-up and surveillance colonoscopies.

## Methods

### Microsimulation model

The outcomes of CRC-AIM have been qualitatively and quantitatively validated against the CISNET models.^14^ Full details of the model have been described elsewhere.^14^ As with all CRC microsimulation models, CRC-AIM has natural history and screening components (additional details in **Supplemental Methods)**. The natural history component models the progression from adenoma development, to preclinical cancer, to symptomatic CRC in unscreened patients. The purpose of CRC screening is to reduce cancer deaths and cancer incidence. This is accomplished by detecting preclinical cancers and adenomas and facilitating the removal of any detected adenomas. The effectiveness of a CRC screening modality is dependent on the test performance (e.g. sensitivity, specificity) and patient adherence, which are reflected in the screening component of CRC-AIM. Patient adherence is assumed to be 100%, per established convention.^4^ As with the CISNET models,^4^ it is assumed in CRC-AIM that a follow-up colonoscopy is always conducted after a positive non-invasive screening test. After a negative follow-up colonoscopy, individuals are assumed to return to their original stool-based screening test and the next screening is due 10 years later. After a positive follow-up colonoscopy, repeated colonoscopies (surveillance colonoscopies) are conducted until at least age 85. The frequency of the surveillance colonoscopies is dependent on the findings of the latest colonoscopy.

### Test performance assumptions

The base-case (“no impact”) adenoma sensitivity values were the same as those used by the CISNET models and were identical for screening and combined follow-up and surveillance colonoscopies (follow-up/surveillance).^3,4^ The base-case (scenario 1; “no impact”) sensitivity values were 75% for adenomas 1 to 5 mm (small adenoma), 85% for adenomas 6 to 9 mm (medium adenoma), and 95% for adenomas greater than 10 mm (large adenoma), as previously reported.^3,4^

Due to the uncertainty and variability of real-world colonoscopy performance, a range of adenoma sensitivity scenarios was assessed (**Table 1**). The investigated ranges of adenoma sensitivity values for additional colonoscopy performance scenarios were developed using different slopes of odds ratio (OR) adjustments. The anchor point for the ranges was a conservative assumption that there was no difference in screening versus follow-up/surveillance colonoscopy sensitivity for adenomas greater than 10 mm.^13^ This corresponds to an OR of 1 and a log(OR) of 0. Sensitivity values between screening and follow-up/surveillance colonoscopies for large adenomas for scenarios 2 to 4, scenarios 5 to 7, and scenarios 8 to 10 were fixed with a ln(OR) of 0 (“small impact”), 1.00 (“medium impact”), or 2.00 (“large impact”), respectively, and then assumed a constant increase in slope of 0.15, 0.30, or 0.60 between large and medium adenomas, and between medium and small adenomas (**Table 1**).

**Table 1.**
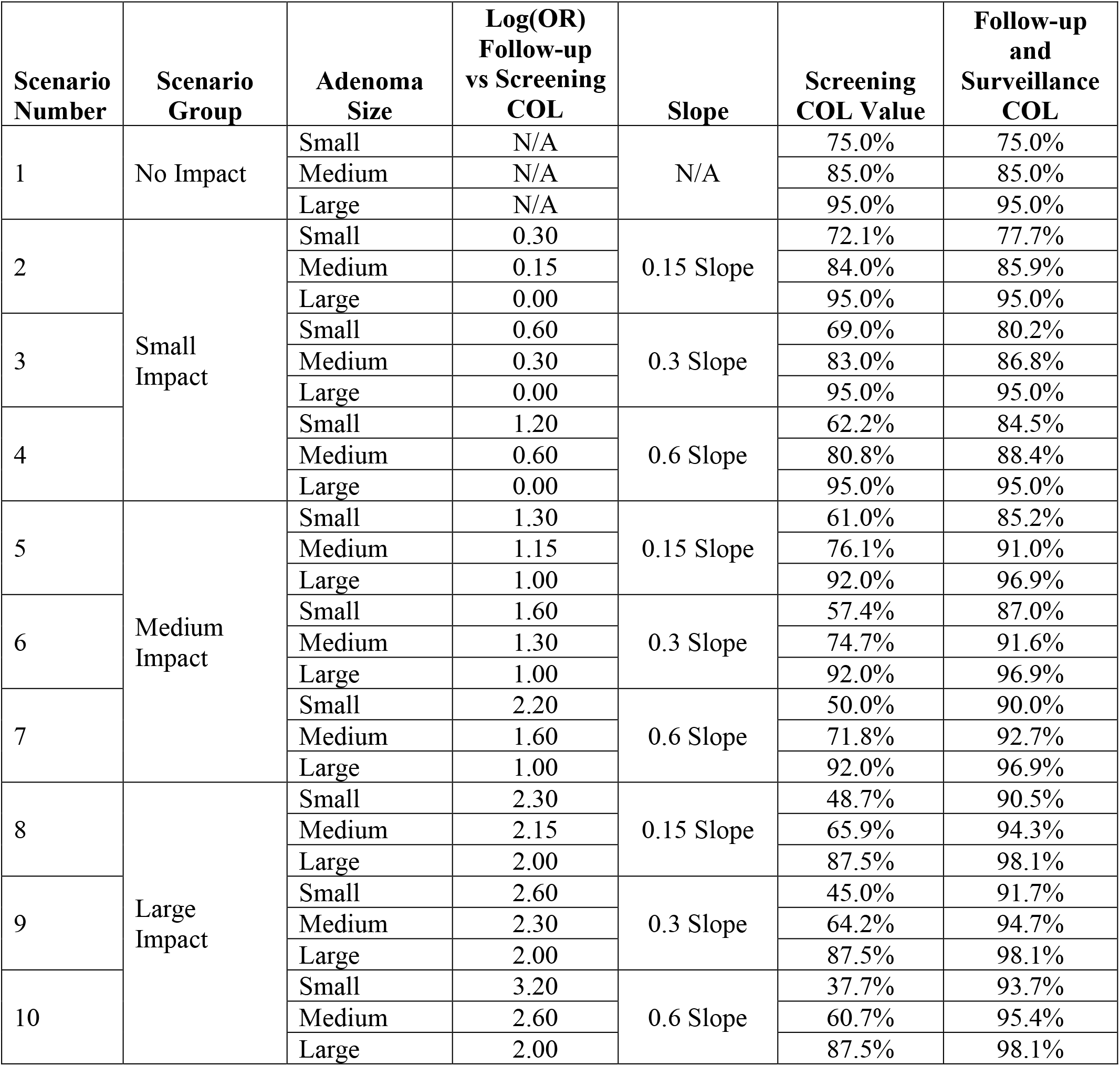
Scenarios for detection sensitivity by adenoma size for screening and follow-up/ surveillance colonoscopy (COL). Base-case (“no impact”) values are those used in CISNET microsimulation models.^3^ Adenomas are defined as small (1–5 mm), medium (6–9 mm), or large (≥10 mm).

In contrast to the colonoscopy adenoma sensitivity scenarios described above, the adenoma sensitivities for FIT and mt-sDNA remained intentionally identical to the CISNET models.^3,4^ The specificity, reach, complications, and sensitivity of CRC detection by disease stage for colonoscopy, FIT, and mt-sDNA were also the same as those used by CISNET models (**Table 2** and **Supplementary Figure S1)**.^3,4^

**Table 2.** Screening test characteristic inputs. Reproduced and adapted with permission from Knudsen et al, 2016.^3^

| Screening test characteristic | Screening Test, Source Citation for Input |  |  |
| --- | --- | --- | --- |
|  | Colonoscopy (within reach, per lesion) | FIT (per person) | mt-sDNA (per person) |
| Sensitivity for CRC, % | 95<sup>a</sup> | 73.8<sup>18</sup> | 92.3<sup>18</sup> |
| Sensitivity for adenomas $\geq 10$ mm, % | Variable by analysis scenario | 23.8<sup>b, 18</sup> | 42.4<sup>b, 18</sup> |
| Sensitivity for adenomas 6-9 mm, % | Variable by analysis scenario | 7.6<sup>c, 18</sup> | 17.2<sup>c, 18</sup> |
| Sensitivity for adenomas 1-5 mm, % | Variable by analysis scenario |  |  |
| Specificity, % | 86<sup>d, 25</sup> | 96.4<sup>18</sup> | 89.8<sup>18</sup> |
| Reach, % | 95 to end of cecum, remainder between rectum and cecum<sup>a,e</sup> | Whole colorectum<sup>a</sup> | Whole colorectum<sup>a</sup> |
| Risk of complications (serious GI, other GI, and CV complications) | Age-specific risks<sup>f, 26-28</sup> | 0<sup>29</sup> | 0<sup>29</sup> |

COL, colonoscopy; CRC, colorectal cancer; CV, cardiovascular; FIT, fecal immunochemical test with a positivity cutoff of $\geq 100$ ng of hemoglobin (Hb) per mL of buffer ( $\geq 20$ mcg Hb/g of feces); GI, gastrointestinal; mt-sDNA, multitarget stool DNA test.
<sup>a</sup>No source, input is by assumption.
<sup>b</sup>Sensitivity for persons with advanced adenomas (ie, adenomas $\geq 10$ mm or adenomas with advanced histology). Sensitivity was not reported for the subset of persons with $\geq 10$ mm adenomas.
<sup>c</sup>Sensitivity for persons with nonadvanced adenomas.
<sup>d</sup>The lack of specificity with endoscopy reflects the detection of nonadenomatous polyps, which, in the case of colonoscopy, leads to unnecessary polypectomy, which is associated with an increased risk of colonoscopy complications.
<sup>e</sup>Full reach (to the cecum) is assumed to be achieved 95% of the time and if reach is only partial, a second colonoscopy is performed.<sup>3</sup>
<sup>f</sup>See **Fig S1** in the Supplement for details on age-specific risks.

### Screening strategies

Intervals for screening used in the model were colonoscopy every 10 years, FIT every 1 year, and mt-sDNA every 3 years. The average-risk screening period was modeled between 50–75 years of age. These intervals and the age range were selected since they are “strong” recommendations from USPSTF and the ACS.^1,3^

### CRC screening outcomes

Simulated outcomes included the numbers of stool tests, complications from colonoscopies, CRC cases, CRC deaths, and life-years with CRC, as well as the number of life-years gained (LYG) and reduction in CRC-related incidence and mortality compared with no CRC screening. Number of colonoscopies was used as an indicator of related resource use, costs, and complications. By virtue of simulated model outputs, AMR (proportion of missed adenomas per colonoscopy) and ADR (proportion of individuals with detected adenomas at a given age) were able to be calculated. Adenoma miss rates have been estimated by tandem colonoscopies in research settings and may be as high as 47.9%.^15^

The weighted mean AMR was calculated using the cross-sectional AMR per colonoscopy. The ADR for FIT and mt-sDNA was calculated for the first follow-up colonoscopy. The percentage of LYG relative to colonoscopy was determined for mt-sDNA and FIT; strategies with LYG within 90% are considered to have comparable effectiveness to colonoscopy.^1,3,4^ All outcomes were simulated for 4 million individuals born in 1975 and were reported per 1,000 individuals free of diagnosed CRC at age 40.

### Sensitivity analysis

Two sensitivity analyses were conducted. In the primary analysis, it was assumed that for each scenario (except the base-case scenario 1) the follow-up/surveillance colonoscopy after a positive mt-sDNA or FIT had a greater sensitivity than a screening colonoscopy based on data published by Johnson et al,^13^ which found that endoscopists who were aware of a positive mt-sDNA found significantly more adenomas when compared with those endoscopists who were not aware of a positive mt-sDNA. However, the study only evaluated mt-sDNA and no similar data are available for FIT. Thus, in the first sensitivity analysis, the primary analysis was replicated with the assumption that the sensitivity of a follow-up/surveillance colonoscopy after FIT was the same as a screening colonoscopy.

For the second sensitivity analysis, the goal was to use the most up-to-date evidence of screening modality performance. The primary analysis was replicated using newer data^16^ for colonoscopy sensitivity combined with more detailed FIT and mt-sDNA sensitivity and age-related specificity values. In the CISNET models^3,4^ and the primary analysis, the base-case (“no impact”) sensitivity values by adenoma size for colonoscopy were identical for screening and follow-up/surveillance and were based on a 2006 meta-analysis of AMR from 6 tandem colonoscopy studies in a total of 465 patients.^17^ Updated base-case detection sensitivity values using AMRs from a 2019 meta-analysis of 44 studies and more than 15,000 tandem colonoscopies were applied to the colonoscopy performance scenarios (**Supplemental Table S1**).^16^ Since there are not high-quality data to support an improved detection rate of follow-up/surveillance colonoscopy versus screening colonoscopy for large adenomas, identical sensitivity values were used for large adenomas (≥10 mm).

The test performance characteristics for FIT and mt-sDNA used in the CISNET models and the primary analysis were derived from data generated in a cross-sectional study (DeeP-C study; clinicaltrials.gov identifier, NCT01397747).^18^ The published report of the cross-sectional study did not distinguish adenomas by size or location, but rather as advanced (≥10 mm) or non-advanced adenomas.^18^ Therefore, the sensitivity of advanced adenomas was used in the primary analysis as a proxy for the sensitivity of adenomas greater than 10 mm and the sensitivity of non-advanced adenomas was used as a proxy for the sensitivity of adenomas 1 to 5 mm and 6 to 9 mm combined.^4^ More detailed estimates of sensitivity by adenoma size and location (rectal, distal, proximal), and age-based specificity, were derived for FIT and mt-sDNA using data collected in the cross-sectional study^18^ to which the study sponsor had access (**Supplemental Table S2**).

All other modeling aspects in the sensitivity analyses were the same as the primary analyses.

## Results

### CRC-related outcomes

For the base-case scenario (scenario 1), the LYG was higher for colonoscopy every 10 years (351.9) compared with triennial mt-sDNA (299.5) and annual FIT (317.8; **Figure 1**). The percentage of LYG relative to colonoscopy was 85% for mt-sDNA and 90% for FIT (**Figure 2**). The reductions in CRC-related incidence and mortality were higher for colonoscopy (83.1% and 85.7%, respectively) compared with mt-sDNA (64.5% and 72.2%) and FIT (68.3% and 76.2%; **Figure 1**). The total number of colonoscopies associated with the every 10 year colonoscopy screening strategy was more than double (4,167) that of triennial mt-sDNA (1,958) or annual FIT (2,036; **Supplemental Table S3**).

**Figure 1.**
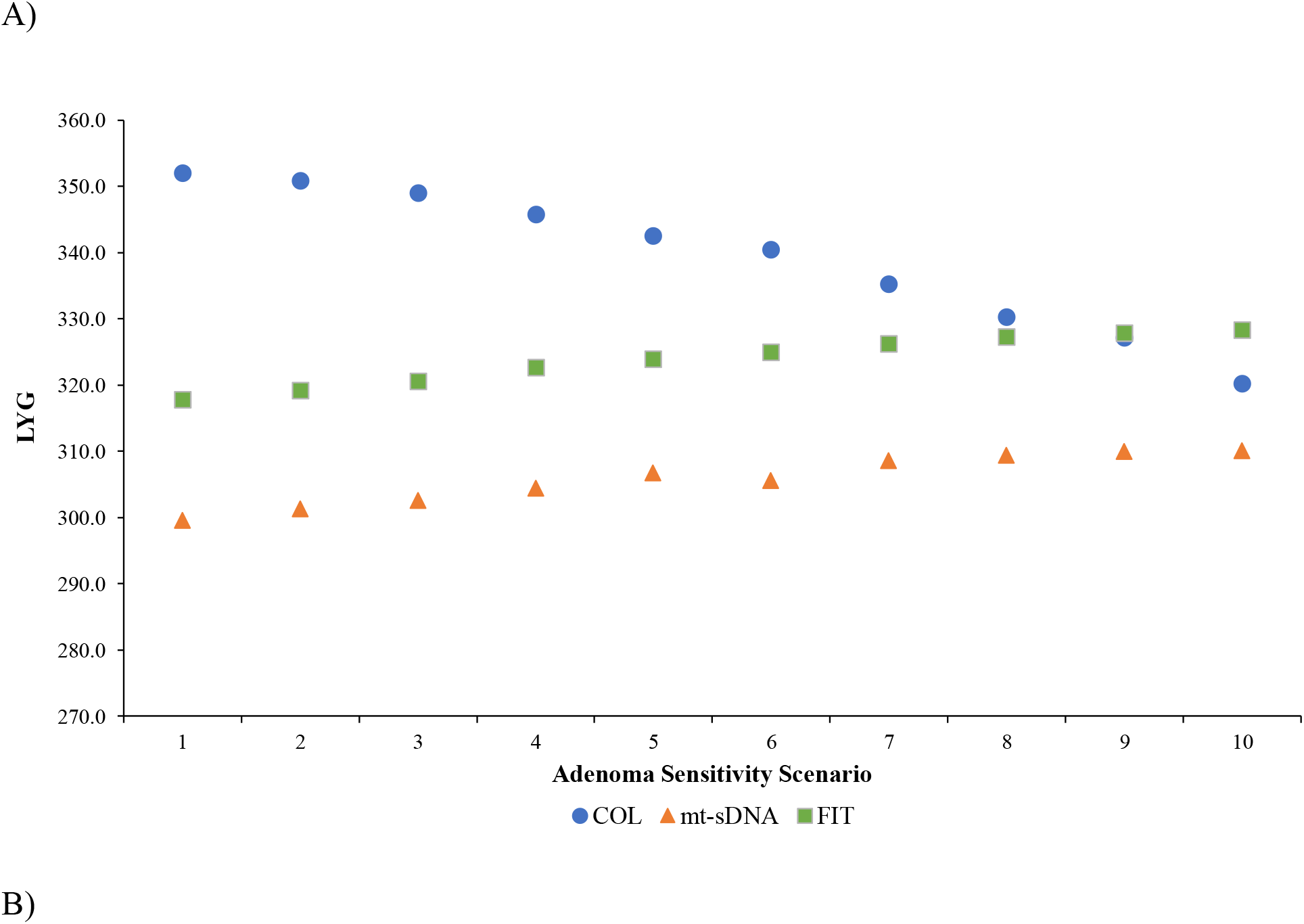
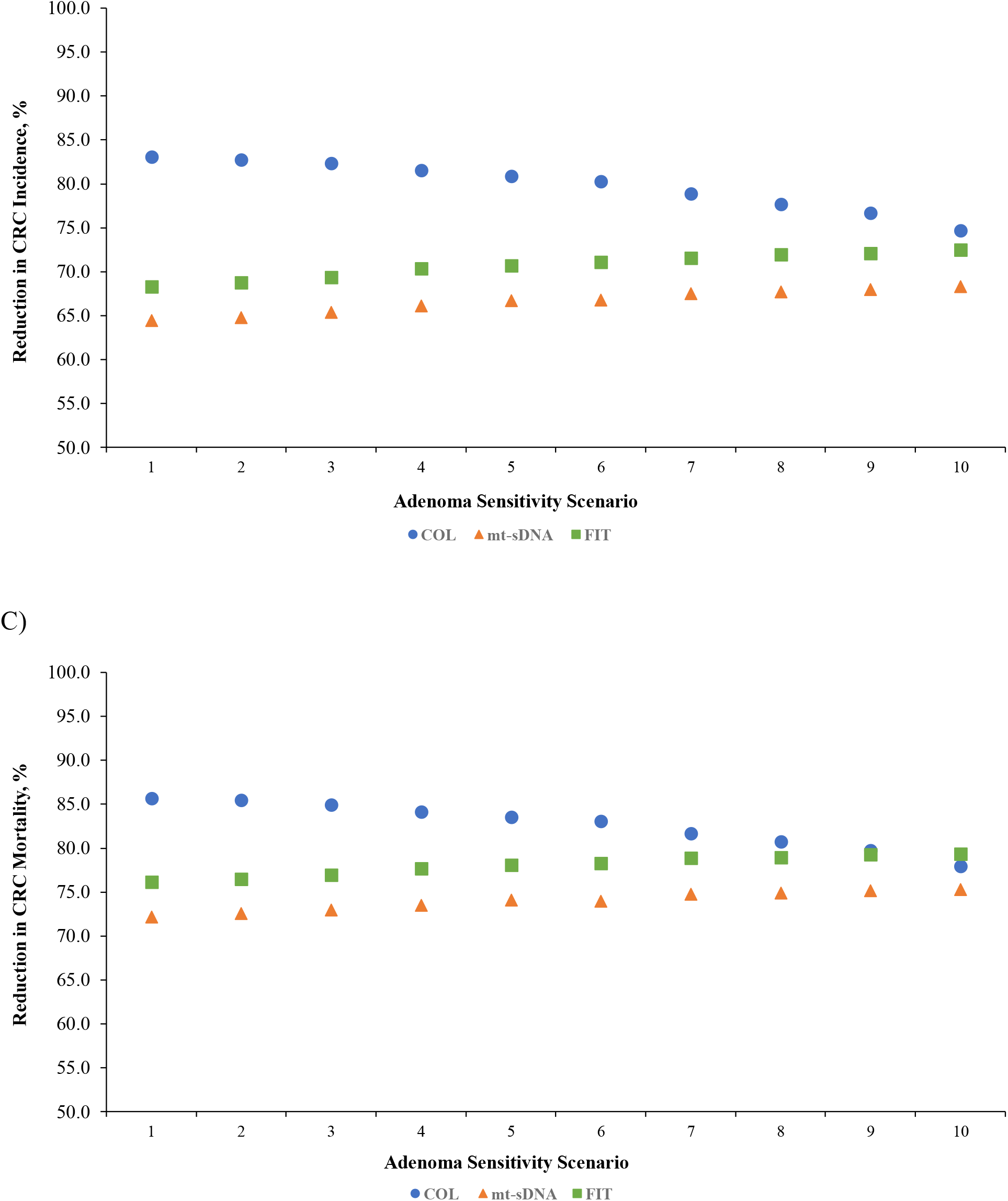
A) Predicted life-years gained (LYG), B) reduction in CRC-related incidence, and C) reduction in CRC-related mortality in scenarios of screening and follow-up/surveillance colonoscopy (COL) adenoma sensitivity. Results are per 1000 individuals screened with COL every 10 years, multitarget stool DNA test (mt-sDNA) every 3 years, or fecal immunochemical test (FIT) every 1 year from ages 50–75 compared with no screening.

**Figure 2.**
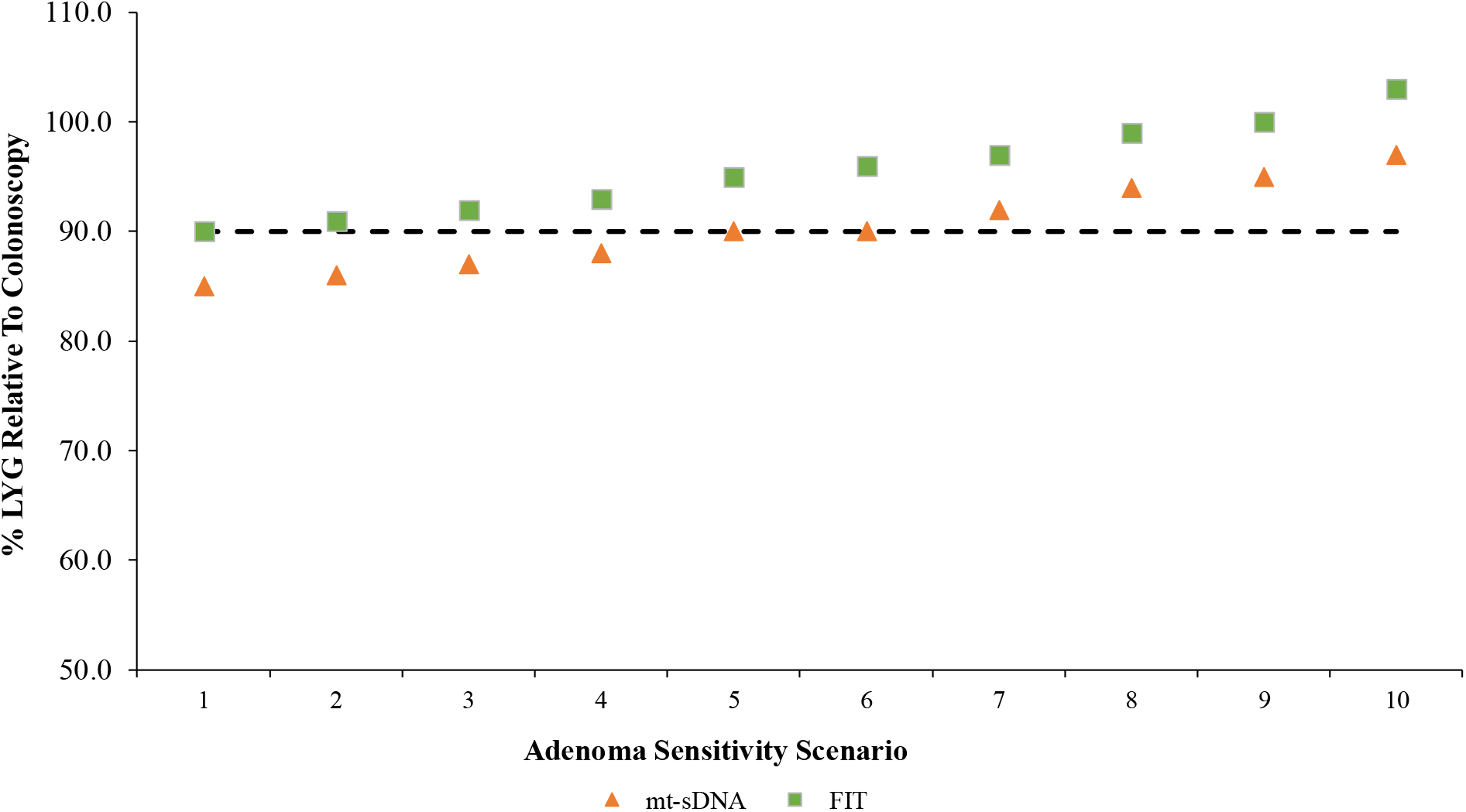
Percentage of predicted life-years gained (LYG) for multitarget stool DNA test (mt-sDNA) and fecal immunochemical test (FIT) relative to LYG by colonoscopy. Data are from different scenarios of screening and follow-up/surveillance colonoscopy adenoma sensitivity. Results are per 1000 individuals screened with mt-sDNA every 3 years or FIT every 1 year from ages 50–75 compared with no screening. The dashed line indicates the 90% LYG threshold which signals comparative effectiveness to colonoscopy.

As the modeled differences in adenoma sensitivity between screening and follow-up/surveillance colonoscopies increased, the LYG and reductions in CRC-related incidence and mortality with colonoscopy every 10 years decreased (**Figure 1**). Compared with the base-case scenario 1, by scenario 10 the LYG with colonoscopy had decreased by 31.7, to 320.2 LYG and the reductions in CRC-related incidence and mortality decreased approximately 8% to 74.7% and 78.0%, respectively (**Supplemental Table S3**). As the modeled differences in adenoma sensitivity increased, the LYG and reductions in CRC-related incidence and mortality with triennial mt-sDNA and annual FIT improved (**Figure 1**). Compared with the base-case scenario, by scenario 10 the LYG with mt-sDNA had increased by 10.5 to 310.0 LYG, and the reductions in CRC-related incidence and mortality increased approximately 3% to 4% to 68.3% and 75.3%, respectively; the LYG with FIT increased by 10.5 LYG to 328.3 LYG, and the reductions in CRC-related incidence and mortality increased approximately 3% to 4% to 72.5% and 79.4%, respectively (**Supplemental Table S3**). The percentage of LYG for mt-sDNA and FIT relative to colonoscopy increased with each scenario (**Figure 2**). The 90% threshold of LYG relative to colonoscopy was reached by FIT in scenario 1, the base-case, and by mt-sDNA at scenario 5, which was defined as one of the “medium impact” scenarios.

### Adenoma miss rates

For the base-case scenario (scenario 1), the weighted mean AMR for colonoscopy every 10 years (21.3%) was greater than that of triennial mt-sDNA (18.9%) and annual FIT (19.0%; **Figure 3A**). As the modeled differences in adenoma sensitivity between screening and follow-up/surveillance colonoscopies increased, the AMR with colonoscopy every 10 years increased until reaching 52.7% at scenario 10. The AMR with triennial mt-sDNA and annual FIT decreased as the modeled differences in adenoma sensitivity increased (**Figure 3A**). By scenario 10, the AMR reached 5.1% for triennial mt-sDNA and 5.1% for annual FIT.

**Figure 3.**
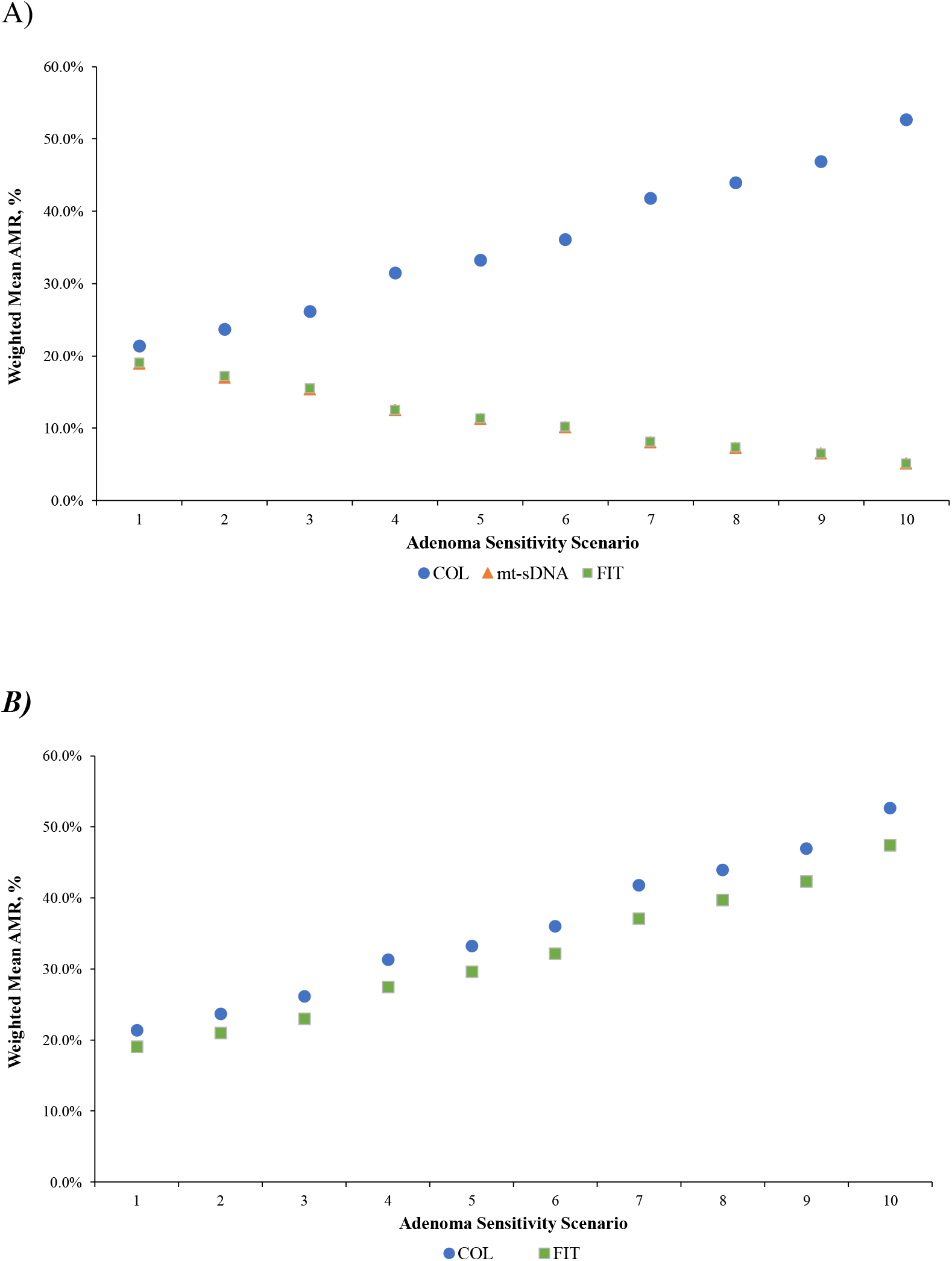
Weighted average adenoma miss rates (AMR) in A) the primary analysis and B) in sensitivity analysis when screening colonoscopy sensitivity is assumed to be the same as a follow-up/surveillance colonoscopy after a positive FIT. Scenarios are of screening and follow-up/surveillance COL adenoma sensitivities.

### Adenoma detection rates

Adenoma detection rates were calculated for the first follow-up colonoscopy after a positive stool-based test (mean age=58.9 y for mt-sDNA and 59.2 y for FIT). For the base-case scenario (scenario 1), the ADR was 30.3% for triennial mt-sDNA and 31.7% for annual FIT (**Supplemental Table S4**). As the modeled differences in adenoma sensitivity between screening and follow-up/surveillance colonoscopies increased, the ADR increased. By scenario 10, the ADR was 33.8% for triennial mt-sDNA and 35.7% for annual FIT.

### Sensitivity analysis

In the first sensitivity analysis that assumed the sensitivity of a follow-up colonoscopy after positive FIT was the same as a screening colonoscopy, the pattern of changes in CRC-related outcomes, AMR, and ADR for FIT over the spectrum of adenoma sensitivity scenarios reversed from that of the primary analysis (**Figure 3B**, **Figure 4**, **Supplemental Table S4**, and **Supplemental Table S5**). In the second sensitivity analysis, the overall pattern of changes in CRC-related outcomes and AMR over the spectrum of adenoma sensitivity scenarios were generally similar in the sensitivity analysis using updated test performance inputs compared with the primary analysis (**Supplemental Figure S2, Supplemental Figure S3,** and **Supplemental Table S6**).

**Figure 4.**
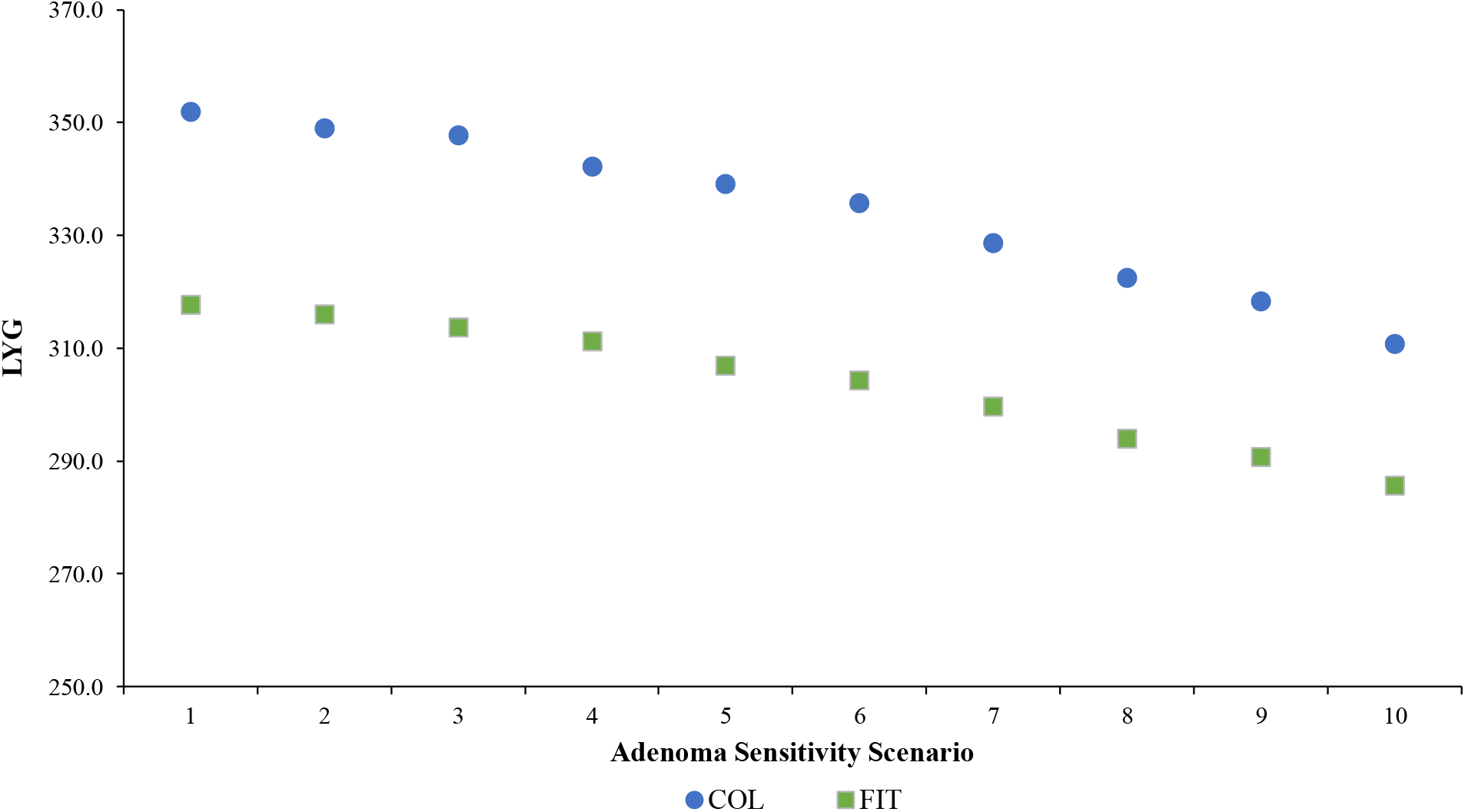
Predicted life-years gained (LYG) in sensitivity analysis when screening colonoscopy sensitivity is assumed to be the same as a follow-up colonoscopy after a positive FIT. Data are from different scenarios of screening and follow-up/surveillance colonoscopy adenoma sensitivity. Results are per 1000 individuals screened with COL every 10 years or fecal immunochemical test (FIT) every 1 year from ages 50–75 compared with no screening.

## Discussion

The results of this CRC-AIM microsimulation study demonstrate the impact of differences in adenoma sensitivity for screening colonoscopy and follow-up/surveillance colonoscopy after a positive stool-based test with respect to multiple CRC outcomes, including AMR and ADR. Within the range of simulated scenarios, when assuming identical adenoma sensitivities regardless of colonoscopy indication, the screening strategy of colonoscopy every 10 years yielded increases in LYG, reductions in CRC incidence and mortality, and higher AMR compared with triennial mt-sDNA and annual FIT. When real-world data were applied to the adenoma sensitivity inputs,^13^ the predicted outcomes progressively improved for the stool-based tests compared with the base-case scenario, particularly as the modeled differences in sensitivities between screening and follow-up/surveillance colonoscopies increased. These results indicate that previous CRC screening microsimulation analyses assuming identical adenoma sensitivities for screening and follow-up colonoscopies artificially underestimated the benefits of stool-based tests and overestimated the benefits of colonoscopy.^3^ Subsequently, FIT was shown to reach the 90% LYG threshold used as a marker of comparable effectiveness to colonoscopy^1,3,4^ under the base case scenario and mt-sDNA was shown to achieve this threshold under one of the “medium impact” adenoma sensitivity scenarios.

Notably, although the blinded vs unblinded endoscopist study by Johnson et al^13^ was only evaluated for mt-sDNA, the same assumption of increased sensitivity with a follow-up/surveillance colonoscopy was applied to FIT in the current primary analysis. In a sensitivity analysis where this assumption was no longer applied, there was no improvement in predicted outcomes with FIT across the adenoma sensitivity scenarios.

AMR is, in part, a reflection of the sensitivity of a colonoscopy whereas ADR is an established indicator of the colonoscopy quality. In the current model, the estimated AMR for screening colonoscopies ranged from 21% in the base-case adenoma sensitivity scenario to 53% in the highest impact adenoma sensitivity scenario. In contrast, the AMR for colonoscopy after a positive stool-based test decreased from 19% in the base-case scenario to 5% in the highest impact scenario. Although these predicted AMRs span a broad range, they are in line with those published from tandem colonoscopies in clinical trials, which in one review ranged from 47.9% to 5.1%.^19^ A systematic review of studies generally conducted under conditions similar to the assumptions in the current model (i.e, optimal bowel preparations, whole colon) determined an average AMR of 22%.^17^ Similarly, the calculated ADRs in the current model were consistent with tandem colonoscopy studies conducted in patients aged 59 to 60 years in which the ADRs ranged from 24.6% to 27.1%.^20,21^

At present, it remains unknown which of the adenoma sensitivity scenarios is most representative of real-world clinical practice in diverse settings. In the Johnson et al^13^ study, the unblinded endoscopists detected relatively 32% more adenomas than the blinded endoscopists. In the current study, when the ADRs were compared between the primary analysis and first sensitivity analyses across the adenoma sensitivity scenarios, a “medium impact” scenario (scenario 7) was the point at which the relative difference in ADR reached approximately 32% for mt-sDNA. Therefore, in terms of ADR, the “medium impact” adenoma sensitivity scenarios may be the most reflective of real-world practice colonoscopy sensitivities.

A limitation of the current analysis is there is little published data on the adenoma sensitivity of surveillance colonoscopy after a positive stool-test compared with screening colonoscopy. Therefore, it was assumed that surveillance colonoscopy adenoma sensitivity was identical to follow-up colonoscopy sensitivity for our modeled scenarios. Similar to previous CISNET models, the current analysis using the CRC-AIM model is limited in that it does not account for serrated polyps which may account for up to 30% of CRC.^22,23^ In addition, 100% adherence to CRC screening was assumed; previous results from the CRC-AIM model indicate that changing adherence assumptions to reflect more real-world practice has an impact on predicted outcomes.^24^

## Conclusion

Application of more realistic, indication-associated estimates for adenoma sensitivity at screening versus follow-up/surveillance colonoscopy provides more accurate simulation of the CRC screening benefits from primary stool-based screening strategies. Future CRC screening microsimulation models should consider incorporating a range of different sensitivities between screening and follow-up/surveillance colonoscopies.

## Acknowledgements

The authors gratefully acknowledge Andrew Piscitello for his valuable contributions to the methodology development. Medical writing and editorial assistance were provided by Erin P. Scott, PhD, of Maple Health Group, LLC, funded by Exact Sciences Corporation.

## References

1. Wolf AMD, Fontham ETH, Church TR, et al. Colorectal cancer screening for average-risk adults: 2018 guideline update from the American Cancer Society. CA Cancer J Clin. 2018;68(4):250–281.

2. Bibbins-Domingo K, Grossman DC, Curry SJ, et al. Screening for Colorectal Cancer: US Preventive Services Task Force Recommendation Statement. JAMA. 2016;315(23):2564–2575.

3. Knudsen AB, Zauber AG, Rutter CM, et al. Estimation of Benefits, Burden, and Harms of Colorectal Cancer Screening Strategies: Modeling Study for the US Preventive Services Task Force. JAMA. 2016;315(23):2595–2609.

4. Zauber AG, Knudsen AB, Rutter C, Lansdorp-Vogelaar I, Kuntz KM. Evaluating the benefits and harms of colorectal cancer screening strategies: A collaborative modeling approach. Agency for Healthcare Research and Quality. Available at: https://www.uspreventiveservicestaskforce.org/Home/GetFile/1/16540/cisnet-draft-modeling-report/pdf.

5. Cancer Intervention and Surveillance Modeling Network (CISNET) Model Registry. National Cancer Institute. Available at: https://resources.cisnet.cancer.gov/registry. Accessed July 22, 2019.

6. Rutter CM, Knudsen AB, Marsh TL, et al. Validation of Models Used to Inform Colorectal Cancer Screening Guidelines: Accuracy and Implications. Med Decis Making. 2016;36(5):604–614.

7. Rutter CM, Savarino JE. An evidence-based microsimulation model for colorectal cancer: validation and application. Cancer Epidemiol Biomarkers Prev. 2010;19(8):1992–2002.

8. Loeve F, Boer R, van Oortmarssen GJ, van Ballegooijen M, Habbema JD. The MISCAN-COLON simulation model for the evaluation of colorectal cancer screening. Comput Biomed Res. 1999;32(1):13–33.

9. Cubiella J, Castells A, Andreu M, et al. Correlation between adenoma detection rate in colonoscopy- and fecal immunochemical testing-based colorectal cancer screening programs. United European Gastroenterol J. 2017;5(2):255–260.

10. Hilsden RJ, Bridges R, Dube C, et al. Defining Benchmarks for Adenoma Detection Rate and Adenomas Per Colonoscopy in Patients Undergoing Colonoscopy Due to a Positive Fecal Immunochemical Test. Am J Gastroenterol. 2016;111(12):1743–1749.

11. Kligman E, Li W, Eckert GJ, Kahi C. Adenoma Detection Rate in Asymptomatic Patients with Positive Fecal Immunochemical Tests. Dig Dis Sci. 2018;63(5):1167–1172.

12. Wong JCT, Chiu HM, Kim HS, et al. Adenoma detection rates in colonoscopies for positive fecal immunochemical tests versus direct screening colonoscopies. Gastrointest Endosc. 2019;89(3):607–613.e601.

13. Johnson DH, Kisiel JB, Burger KN, et al. Multitarget stool DNA test: clinical performance and impact on yield and quality of colonoscopy for colorectal cancer screening. Gastrointestinal endoscopy. 2017;85(3):657–665.e651.

14. Piscitello A, Borah B, Matney M, et al. Description and validaton of the novel Colorectal Cancer and Adenoma Incidence & Mortality (CRC-AIM) Microsimulation model. Available at: https://www.biorxiv.org/content/10.1101/2020.03.02.966838v1.

15. Chokshi RV, Hovis CE, Hollander T, Early DS, Wang JS. Prevalence of missed adenomas in patients with inadequate bowel preparation on screening colonoscopy. Gastrointest Endosc. 2012;75(6):1197–1203.

16. Zhao S, Wang S, Pan P, et al. Magnitude, Risk Factors, and Factors Associated With Adenoma Miss Rate of Tandem Colonoscopy: A Systematic Review and Meta-analysis. Gastroenterology. 2019;156(6):1661–1674.e1611.

17. van Rijn JC, Reitsma JB, Stoker J, et al. Polyp miss rate determined by tandem colonoscopy: a systematic review. Am J Gastroenterol. 2006;101(2):343–350.

18. Imperiale TF, Ransohoff DF, Itzkowitz SH, et al. Multitarget stool DNA testing for colorectal-cancer screening. N Engl J Med. 2014;370(14):1287–1297.

19. Wang CL, Huang ZP, Chen K, et al. Adenoma miss rate determined by very shortly repeated colonoscopy: Retrospective analysis of data from a single tertiary medical center in China. Medicine (Baltimore). 2018;97(38):e12297.

20. Chandran S, Parker F, Vaughan R, et al. Right-sided adenoma detection with retroflexion versus forward-view colonoscopy. Gastrointest Endosc. 2015;81(3):608–613.

21. Hewett DG, Rex DK. Miss rate of right-sided colon examination during colonoscopy defined by retroflexion: an observational study. Gastrointest Endosc. 2011;74(2):246–252.

22. Bordaçahar B, Barret M, Terris B, et al. Sessile serrated adenoma: From identification to resection. Dig Liver Dis. 2015;47(2):95–102.

23. Ma MX, Bourke MJ. Sessile Serrated Adenomas: How to Detect, Characterize and Resect. Gut Liver. 2017;11(6):747–760.

24. Piscitello A, Saoud L, Fendrick AM, et al. Estimating the impact of imperfect adherence to stool-based colorectal cancer screening strategies on comparative effectiveness using the CRC-AIM microsimulation model. Gastroenterology. 2020;158 (Suppl 1)(6):S–910.

25. Schroy PC, 3rd, Coe A, Chen CA, O’Brien MJ, Heeren TC. Prevalence of advanced colorectal neoplasia in white and black patients undergoing screening colonoscopy in a safety-net hospital. Ann Intern Med. 2013;159(1):13–20.

26. Gatto NM, Frucht H, Sundararajan V, et al. Risk of perforation after colonoscopy and sigmoidoscopy: a population-based study. J Natl Cancer Inst. 2003;95(3):230–236.

27. van Hees F, Zauber AG, Klabunde CN, et al. The appropriateness of more intensive colonoscopy screening than recommended in Medicare beneficiaries: a modeling study. JAMA Intern Med. 2014;174(10):1568–1576.

28. Warren JL, Klabunde CN, Mariotto AB, et al. Adverse events after outpatient colonoscopy in the Medicare population. Ann Intern Med. 2009;150(12):849–857, W152.

29. Lin JS, Piper MA, Perdue LA, et al. Screening for Colorectal Cancer: A Systematic Review for the U.S. Preventive Services Task Force. Evidence Synthesis No. 135. AHRQ Publication No. 14-05203-EF-1. Rockville (MD): Agency for Healthcare Research and Quality; 2016.

