## Supplemental material for "Impact of Screening and Follow-up Colonoscopy Adenoma Sensitivity on Colorectal Cancer Screening Outcomes in the CRC-AIM Microsimulation Model"

**Methods**

*Microsimulation model*

A non-homogenous Poisson process based on sex, age, and person-specific risk is used to generate adenomas within an individual and the generated adenomas are assigned a location in either the colon or rectum.[1,2] The growth rate of adenomas, based on location, are assumed to follow a non-linear growth curve. The cumulative probability of an adenoma transitioning to preclinical CRC is assumed to be a function of adenoma size and location, age at adenoma initiation, and sex. Once the transition occurs, the preclinical CRC is initially assigned a size of 0.5 mm, the sojourn time is determined, and the size of the preclinical CRC upon reaching sojourn time is sampled from a Surveillance, Epidemiology, and End Results (SEER) distribution of CRC sizes. Simple exponential growth of the CRC is assumed. The CRC size is used to determine the cancer stage at diagnosis and the cancer stage at detection is further determined using a multinomial logistic regression model.[1,2] All-cause (non-CRC) mortality rates are from sex-specific cohort life tables from the years 1900-2010. The survival based on CRC-stage is from parametric regression models developed from SEER data for cancers diagnosed from 2000-2003.

The screening component of the model assumes that CRC screening enables the detection of adenomas and preclinical lesions and facilitates in the removal of adenomas.[3,4] The effectiveness of the screening test is influenced by adherence, the sensitivity and specificity of the screening test, and for colonoscopy, the reach of the test. Sensitivity inputs for mt-sDNA and FIT are per person and are based on the characteristics of the most advanced lesion. Sensitivity inputs for colonoscopy are per lesion. False positives are possible. Complications of serious and nonserious gastrointestinal events or cardiovascular events resulting from adenoma removal can occur and are factored into the screening component.

### References


1. CISNET Colorectal Cancer Collaborators. RAND Corporation (CRC-SPIN), 2015. HI.001.03112015.70373. National Cancer Institute Cancer Intervention and Surveillance Modeling Network. Available at: https://cisnet.cancer.gov/colorectal/profiles.html. Accessed November 21, 2019.
2. Rutter CM, Miglioretti DL, Savarino JE. Bayesian Calibration of Microsimulation Models. *J Am Stat Assoc.* 2009;104(488):1338-1350.
3. Knudsen AB, Zauber AG, Rutter CM, et al. Estimation of Benefits, Burden, and Harms of Colorectal Cancer Screening Strategies: Modeling Study for the US Preventive Services Task Force. *JAMA.* 2016;315(23):2595-2609.
4. Zauber AG, Knudsen AB, Rutter C, Lansdorp-Vogelaar I, Kuntz KM. Evaluating the benefits and harms of colorectal cancer screening strategies: A collaborative modeling approach. Agency for Healthcare Research and Quality. Available at: https://www.uspreventiveservicestaskforce.org/Home/GetFile/1/16540/cisnet-draft-modeling-report/pdf.
5. Zhao S, Wang S, Pan P, et al. Magnitude, Risk Factors, and Factors Associated With Adenoma Miss Rate of Tandem Colonoscopy: A Systematic Review and Meta-analysis. *Gastroenterology.* 2019;156(6):1661-1674.e1611.
6. Imperiale TF, Ransohoff DF, Itzkowitz SH, et al. Multitarget stool DNA testing for colorectal-cancer screening. *N Engl J Med.* 2014;370(14):1287-1297.

7. Lin JS, Piper MA, Perdue LA, et al. Screening for Colorectal Cancer: A Systematic Review for the U.S. Preventive Services Task Force. Evidence Synthesis No. 135. AHRQ Publication No. 14-05203-EF-1. Rockville (MD): Agency for Healthcare Research and Quality; 2016.

8. van Hees F, Zauber AG, Klabunde CN, et al. The appropriateness of more intensive colonoscopy screening than recommended in Medicare beneficiaries: a modeling study. *JAMA Intern Med.* 2014;174(10):1568-1576.

9. Warren JL, Klabunde CN, Mariotto AB, et al. Adverse events after outpatient colonoscopy in the Medicare population. *Ann Intern Med.* 2009;150(12):849-857, W152.

**Table S1**. Scenarios for adenoma sensitivity for screening and follow-up/surveillance colonoscopy (COL) using base-case ("no impact") values derived from adenoma miss rate data in a meta-analysis published in 2019.[5] Adenomas are defined as small (1–5 mm), medium (6–9 mm), or large (≥10 mm).

| Scenario Number | Scenario Group | Slope | Adenoma Size | Screening COL Value | Follow-up and Surveillance COL |
|---|---|---|---|---|---|
| 1 | No Impact | N/A | Small | 69.0% | 69.0% |
| | | | Medium | 81.0% | 81.0% |
| | | | Large | 91.0% | 91.0% |
| 2 | Small Impact | 0.15 Slope | Small | 65.7% | 72.1% |
| | | | Medium | 79.8% | 82.1% |
| | | | Large | 91.0% | 91.0% |
| 3 | | 0.3 Slope | Small | 62.2% | 75.0% |
| | | | Medium | 78.6% | 83.2% |
| | | | Large | 91.0% | 91.0% |
| 4 | | 0.6 Slope | Small | 55.0% | 80.2% |
| | | | Medium | 76.0% | 85.2% |
| | | | Large | 91.0% | 91.0% |
| 5 | Medium Impact | 0.15 Slope | Small | 53.7% | 81.0% |
| | | | Medium | 70.6% | 88.3% |
| | | | Large | 86.0% | 94.3% |
| 6 | | 0.3 Slope | Small | 50.0% | 83.2% |
| | | | Medium | 69.0% | 89.1% |
| | | | Large | 86.0% | 94.3% |
| 7 | | 0.6 Slope | Small | 42.6% | 87.0% |
| | | | Medium | 65.7% | 90.5% |
| | | | Large | 86.0% | 94.3% |

| | | | | | |
|---|---|---|---|---|---|
| 8 | Large Impact | 0.15 Slope | Small | 41.3% | 87.5% |
| | | | Medium | 59.3% | 92.6% |
| | | | Large | 78.8% | 96.5% |
| 9 | | 0.3 Slope | Small | 37.8% | 89.1% |
| | | | Medium | 57.4% | 93.1% |
| | | | Large | 78.8% | 96.5% |
| 10 | | 0.6 Slope | Small | 31.0% | 91.7% |
| | | | Medium | 53.7% | 94.0% |
| | | | Large | 78.8% | 96.5% |

**Table S2**. FIT and mt-sDNA screening characteristics for sensitivity analyses. Adenoma sensitivity values by adenoma size and location and specificity by age were derived from a cross-sectional study (clinicaltrials.gov identifier, NCT01397747).[6]

| | **Sensitivity** | | | | | | | | | | **Specificity** | | | | |
|---|---|---|---|---|---|---|---|---|---|---|---|---|---|---|---|
| | **Adenomas, mm** | | | | | | | | | | | | | | |
| | **Rectal** | | | **Distal** | | | **Proximal** | | | **Cancer Stage** | **Age, y** | | | | |
| | **≤5** | **6-9** | **≥10** | **≤5** | **6-9** | **≥10** | **≤5** | **6-9** | **≥10** | **I-IV** | **<60** | **60-64** | **65-69** | **70-74** | **75+** |
| FIT, % (n/N) | 3.3% (5/150) | 5.6% (5/89) | 30.1% (22/73) | 7.0% (39/558) | 21.5% (68/317) | 39.3% (83/211) | 5.9% (73/1236) | 6.6% (40/609) | 16.0% (65/406) | 73.8% (48/65) | 97.8% (1361/1392) | 96.5% (355/368) | 95.6% (1483/1551) | 96.7% (697/721) | 93.9% (399/425) |
| 95% CI | 1.1% - 7.6% | 1.8% - 12.6% | 19.9% - 42.0% | 5.0% - 9.4% | 17.1% - 26.4% | 32.7% - 46.3% | 4.7% - 7.4% | 4.7% - 8.8% | 12.6% - 19.9% | 61.5% - 84.0% | 96.9% - 98.5% | 94.0% - 98.1% | 94.5% - 96.6% | 95.1% - 97.9% | 91.2% - 96.0% |
| mt-sDNA, % (n/N) | 9.3% (14/150) | 21.3% (19/89) | 56.2% (41/73) | 14.5% (81/558) | 27.8% (88/317) | 57.3% (121/211) | 15.8% (195/1236) | 19.9% (121/609) | 34.0% (138/406) | 92.3% (60/65) | 94.4% (1314/1392) | 92.4% (340/368) | 89.1% (1382/1551) | 86.1% (621/721) | 81.2% (345/425) |
| 95% CI | 5.2% - 15.2% | 13.4% - 31.3% | 44.1% - 67.8% | 11.7% - 17.7% | 22.9% - 33.0% | 50.4% - 64.1% | 13.8% - 17.9% | 16.8% - 23.3% | 29.4% - 38.8% | 83.0% - 97.5% | 93.1% - 95.5% | 89.2% - 94.9% | 87.4% - 90.6% | 83.4% - 88.6% | 77.1% - 84.8% |

FIT, fecal immunochemical test; mt-sDNA, multitarget stool DNA test.

2-sided 95% confidence intervals calculated using the Exact (Clopper-Pearson) method.

**Table S3.** Screening outcomes in scenarios of screening and follow-up/surveillance COL adenoma sensitivity. Results are per 1000 individuals screened with COL every 10 years, mt-sDNA every 3 years, or FIT every 1 year from ages 50–75 compared with no screening.

| Colonoscopy Performance Scenario | Screening Strategy | Stool Tests | Follow-up COLs | Surveillance COLs | Total COLs | Complications | CRC Cases | CRC Deaths | LY with CRC | LYG | Incidence Reduction | Mortality Reduction | % Threshold LYG Relative to COL | AMR, Weighted Mean |
|---|---|---|---|---|---|---|---|---|---|---|---|---|---|---|
| | No screening | 0 | 0 | 0 | 80 | 2 | 80.3 | 36.6 | 646.0 | 0.0 | 0.0% | 0.0% | | N/A |
| Scenario 1 | Colonoscopy 50-75, 10 | 0 | 0 | 1,710 | 4,167 | 16 | 13.6 | 5.2 | 154.0 | 351.9 | 83.1% | 85.7% | | 21.3% |
| | mt-sDNA, 50-75, 3 | 6,075 | 813 | 1,128 | 1,958 | 11 | 28.5 | 10.2 | 331.7 | 299.5 | 64.5% | 72.2% | 85% | 18.9% |
| | FIT, 50-75, 1 | 15,857 | 809 | 1,213 | 2,036 | 11 | 25.5 | 8.7 | 309.8 | 317.8 | 68.3% | 76.2% | 90% | 19.0% |
| Scenario 2 | Colonoscopy 50-75, 10 | 0 | 0 | 1,689 | 4,152 | 15 | 13.8 | 5.3 | 156.3 | 350.8 | 82.8% | 85.5% | | 23.7% |
| | mt-sDNA, 50-75, 3 | 6,068 | 813 | 1,140 | 1,970 | 11 | 28.2 | 10.0 | 330.1 | 301.2 | 64.8% | 72.6% | 86% | 17.0% |
| | FIT, 50-75, 1 | 15,836 | 808 | 1,227 | 2,050 | 11 | 25.1 | 8.6 | 306.0 | 319.1 | 68.8% | 76.5% | 91% | 17.2% |
| Scenario 3 | Colonoscopy 50-75, 10 | 0 | 0 | 1,666 | 4,137 | 15 | 14.1 | 5.5 | 159.8 | 349.0 | 82.4% | 85.0% | | 26.1% |
| | mt-sDNA, 50-75, 3 | 6,064 | 810 | 1,151 | 1,978 | 11 | 27.8 | 9.9 | 326.8 | 302.5 | 65.4% | 73.0% | 87% | 15.4% |
| | FIT, 50-75, 1 | 15,847 | 804 | 1,238 | 2,057 | 12 | 24.6 | 8.4 | 303.5 | 320.5 | 69.4% | 77.0% | 92% | 15.5% |
| Scenario 4 | Colonoscopy 50-75, 10 | 0 | 0 | 1,614 | 4,103 | 15 | 14.8 | 5.8 | 165.6 | 345.8 | 81.6% | 84.2% | | 31.4% |
| | mt-sDNA, 50-75, 3 | 6,051 | 809 | 1,171 | 1,997 | 11 | 27.2 | 9.7 | 321.5 | 304.4 | 66.1% | 73.5% | 88% | 12.5% |
| | FIT, 50-75, 1 | 15,797 | 803 | 1,264 | 2,081 | 12 | 23.8 | 8.1 | 295.8 | 322.6 | 70.4% | 77.7% | 93% | 12.5% |
| Scenario 5 | Colonoscopy 50-75, 10 | 0 | 0 | 1,595 | 4,090 | 15 | 15.4 | 6.0 | 171.7 | 342.5 | 80.9% | 83.6% | | 33.2% |
| | mt-sDNA, 50-75, 3 | 6,051 | 806 | 1,181 | 2,003 | 11 | 26.7 | 9.5 | 317.3 | 306.7 | 66.7% | 74.1% | 90% | 11.2% |
| | FIT, 50-75, 1 | 15,803 | 800 | 1,272 | 2,086 | 12 | 23.5 | 8.0 | 293.0 | 323.9 | 70.7% | 78.1% | 95% | 11.3% |
| Scenario 6 | Colonoscopy 50-75, 10 | 0 | 0 | 1,565 | 4,071 | 15 | 15.8 | 6.2 | 176.4 | 340.4 | 80.3% | 83.1% | | 36.1% |
| | mt-sDNA, 50-75, 3 | 6,043 | 808 | 1,188 | 2,012 | 11 | 26.6 | 9.5 | 316.3 | 305.5 | 66.8% | 74.0% | 90% | 10.1% |
| | FIT, 50-75, 1 | 15,788 | 800 | 1,279 | 2,094 | 12 | 23.2 | 7.9 | 290.2 | 324.9 | 71.1% | 78.3% | 96% | 10.2% |

| | | | | | | | | | | | | | | |
|---|---|---|---|---|---|---|---|---|---|---|---|---|---|---|
| Scenario 7 | Colonoscopy 50-75, 10 | 0 | 0 | 1,500 | 4,026 | 15 | 17.0 | 6.7 | 185.9 | 335.2 | 78.9% | 81.7% | | 41.8% |
| | mt-sDNA, 50-75, 3 | 6,036 | 805 | 1,204 | 2,025 | 12 | 26.1 | 9.2 | 313.0 | 308.6 | 67.5% | 74.8% | 92% | 8.0% |
| | FIT, 50-75, 1 | 15,777 | 797 | 1,295 | 2,105 | 12 | 22.8 | 7.7 | 287.7 | 326.2 | 71.6% | 78.9% | 97% | 8.1% |
| Scenario 8 | Colonoscopy 50-75, 10 | 0 | 0 | 1,474 | 4,009 | 15 | 17.9 | 7.0 | 196.7 | 330.3 | 77.7% | 80.8% | | 44.0% |
| | mt-sDNA, 50-75, 3 | 6,037 | 804 | 1,207 | 2,027 | 11 | 26.0 | 9.2 | 310.9 | 309.3 | 67.7% | 74.9% | 94% | 7.3% |
| | FIT, 50-75, 1 | 15,769 | 797 | 1,301 | 2,111 | 12 | 22.5 | 7.7 | 284.0 | 327.2 | 72.0% | 79.0% | 99% | 7.3% |
| Scenario 9 | Colonoscopy 50-75, 10 | 0 | 0 | 1,438 | 3,984 | 15 | 18.7 | 7.4 | 203.9 | 327.1 | 76.7% | 79.8% | | 46.9% |
| | mt-sDNA, 50-75, 3 | 6,035 | 803 | 1,214 | 2,033 | 11 | 25.7 | 9.1 | 308.3 | 309.9 | 68.0% | 75.2% | 95% | 6.5% |
| | FIT, 50-75, 1 | 15,755 | 797 | 1,308 | 2,118 | 12 | 22.4 | 7.6 | 284.3 | 327.8 | 72.1% | 79.3% | 100% | 6.5% |
| Scenario 10 | Colonoscopy 50-75, 10 | 0 | 0 | 1,359 | 3,930 | 15 | 20.4 | 8.1 | 217.5 | 320.2 | 74.7% | 78.0% | | 52.7% |
| | mt-sDNA, 50-75, 3 | 6,027 | 803 | 1,224 | 2,043 | 11 | 25.4 | 9.0 | 306.5 | 310.0 | 68.3% | 75.3% | 97% | 5.1% |
| | FIT, 50-75, 1 | 15,756 | 794 | 1,317 | 2,125 | 12 | 22.1 | 7.5 | 281.4 | 328.3 | 72.5% | 79.4% | 103% | 5.1% |

AMR, adenoma miss rate; COL, colonoscopy; CRC, colorectal cancer; FIT, fecal immunochemical test; LY, life-years; LYG, life-years gained; mt-sDNA, multitarget stool DNA test; N/A, not applicable.

**Table S4.** Adenoma detection rates for the first follow-up colonoscopy. The differential colonoscopy adenoma sensitivity column assumes follow-up colonoscopy after a positive mt-sDNA or FIT is more sensitive than screening colonoscopy (primary analysis). The equal colonoscopy adenoma sensitivity column assumes follow-up colonoscopy after a positive mt-sDNA or FIT has the same sensitivity as screening colonoscopy (first sensitivity analysis). Data are for scenarios of screening and follow-up/surveillance COL adenoma sensitivity.

| Colonoscopy Performance Scenario | Screening Strategy | ADR, % Equal COL Adenoma Sensitivity | ADR, % Differential COL Adenoma Sensitivity | Relative Increase in ADR, % |
|---|---|---|---|---|
| Scenario 1 | mt-sDNA, 50-75, 3 | 30.3 | 30.3 | - |
| | FIT, 50-75, 1 | 31.7 | 31.7 | - |
| Scenario 2 | mt-sDNA, 50-75, 3 | 30.2 | 30.7 | 1.6 |
| | FIT, 50-75, 1 | 31.7 | 32.2 | 1.5 |
| Scenario 3 | mt-sDNA, 50-75, 3 | 29.7 | 31.2 | 5.1 |
| | FIT, 50-75, 1 | 31.1 | 32.8 | 5.2 |
| Scenario 4 | mt-sDNA, 50-75, 3 | 29.1 | 31.9 | 9.5 |
| | FIT, 50-75, 1 | 30.5 | 33.6 | 10.3 |
| Scenario 5 | mt-sDNA, 50-75, 3 | 27.8 | 32.3 | 16.1 |
| | FIT, 50-75, 1 | 29.2 | 34.0 | 16.5 |
| Scenario 6 | mt-sDNA, 50-75, 3 | 26.5 | 32.5 | 22.5 |
| | FIT, 50-75, 1 | 27.7 | 34.2 | 23.5 |
| Scenario 7 | mt-sDNA, 50-75, 3 | 24.9 | 33.1 | 32.7 |
| | FIT, 50-75, 1 | 26.1 | 34.8 | 33.6 |
| Scenario 8 | mt-sDNA, 50-75, 3 | 24.3 | 33.2 | 36.7 |
| | FIT, 50-75, 1 | 25.2 | 35.1 | 39.0 |
| Scenario 9 | mt-sDNA, 50-75, 3 | 23.4 | 33.5 | 43.1 |
| | FIT, 50-75, 1 | 24.3 | 35.2 | 45.2 |
| Scenario 10 | mt-sDNA, 50-75, 3 | 21.8 | 33.8 | 55.3 |
| | FIT, 50-75, 1 | 22.5 | 35.7 | 58.7 |

**Table S5.** Screening outcomes in sensitivity analysis where screening colonoscopy is assumed to be the same as a follow-up colonoscopy after a positive FIT. Outcomes are in scenarios of screening and follow-up/surveillance COL adenoma sensitivity. Results are per 1000 individuals screened with COL every 10 years or FIT every 1 year from ages 50–75 compared with no screening.

| Colonoscopy Performance Scenario | Screening Strategy | Stool Tests | Follow-up COLs | Surveillance COLs | Total COLs | Complications | CRC Cases | CRC Deaths | LY with CRC | LYG | Incidence Reduction | Mortality Reduction | % Threshold LYG Relative to COL | AMR, Weighted Mean |
|---|---|---|---|---|---|---|---|---|---|---|---|---|---|---|
| | No screening | 0 | 0 | 0 | 80 | 2 | 80.3 | 36.6 | 646.0 | 0.0 | 0.0% | 0.0% | | N/A |
| Scenario 1 | Colonoscopy 50-75, 10 | 0 | 0 | 1,710 | 4,167 | 16 | 13.6 | 5.2 | 154.0 | 351.9 | 83.1% | 85.7% | | 21.3% |
| | FIT, 50-75, 1 | 15,857 | 809 | 1,213 | 2,036 | 11 | 25.5 | 8.7 | 309.8 | 317.8 | 68.3% | 76.2% | 90% | 19.0% |
| Scenario 2 | Colonoscopy 50-75, 10 | 0 | 0 | 1,689 | 4,153 | 15 | 14.2 | 5.5 | 159.1 | 349.0 | 82.3% | 85.0% | | 23.6% |
| | FIT, 50-75, 1 | 15,876 | 810 | 1,197 | 2,022 | 11 | 25.9 | 8.9 | 313.2 | 316.0 | 67.7% | 75.6% | 91% | 21.0% |
| Scenario 3 | Colonoscopy 50-75, 10 | 0 | 0 | 1,666 | 4,137 | 15 | 14.8 | 5.7 | 164.1 | 347.7 | 81.6% | 84.5% | | 26.1% |
| | FIT, 50-75, 1 | 15,883 | 812 | 1,181 | 2,009 | 11 | 26.5 | 9.2 | 316.1 | 313.7 | 67.0% | 75.0% | 90% | 23.0% |
| Scenario 4 | Colonoscopy 50-75, 10 | 0 | 0 | 1,613 | 4,101 | 15 | 16.2 | 6.2 | 175.0 | 342.1 | 79.8% | 83.0% | | 31.3% |
| | FIT, 50-75, 1 | 15,926 | 816 | 1,146 | 1,979 | 11 | 27.5 | 9.5 | 325.5 | 311.2 | 65.7% | 74.1% | 91% | 27.5% |
| Scenario 5 | Colonoscopy 50-75, 10 | 0 | 0 | 1,595 | 4,090 | 15 | 17.0 | 6.5 | 184.2 | 339.1 | 78.8% | 82.2% | | 33.2% |
| | FIT, 50-75, 1 | 15,947 | 818 | 1,126 | 1,962 | 11 | 28.7 | 9.9 | 337.2 | 306.9 | 64.3% | 73.1% | 91% | 29.6% |
| Scenario 6 | Colonoscopy 50-75, 10 | 0 | 0 | 1,565 | 4,071 | 14 | 17.9 | 6.8 | 191.7 | 335.7 | 77.8% | 81.4% | | 36.0% |
| | FIT, 50-75, 1 | 15,970 | 821 | 1,105 | 1,943 | 10 | 29.3 | 10.2 | 341.1 | 304.3 | 63.5% | 72.3% | 91% | 32.1% |
| Scenario 7 | Colonoscopy 50-75, 10 | 0 | 0 | 1,499 | 4,025 | 14 | 19.8 | 7.6 | 207.8 | 328.6 | 75.3% | 79.3% | | 41.8% |
| | FIT, 50-75, 1 | 16,023 | 825 | 1,063 | 1,907 | 10 | 31.0 | 10.7 | 355.0 | 299.7 | 61.5% | 70.7% | 91% | 37.1% |
| Scenario 8 | Colonoscopy 50-75, 10 | 0 | 0 | 1,473 | 4,009 | 14 | 21.2 | 8.1 | 221.0 | 322.5 | 73.6% | 77.9% | | 43.9% |
| | FIT, 50-75, 1 | 16,032 | 830 | 1,037 | 1,887 | 10 | 32.3 | 11.3 | 367.8 | 294.0 | 59.8% | 69.2% | 91% | 39.7% |

| | | | | | | | | | | | | | | |
|---|---|---|---|---|---|---|---|---|---|---|---|---|---|---|
| Scenario 9 | Colonoscopy 50-75, 10 | 0 | 0 | 1,435 | 3,983 | 14 | 22.3 | 8.5 | 231.7 | 318.4 | 72.2% | 76.8% | | 46.9% |
| | FIT, 50-75, 1 | 16,060 | 833 | 1,013 | 1,866 | 10 | 33.3 | 11.6 | 376.8 | 290.7 | 58.5% | 68.2% | 91% | 42.3% |
| Scenario 10 | Colonoscopy 50-75, 10 | 0 | 0 | 1,359 | 3,932 | 13 | 24.7 | 9.4 | 250.9 | 310.8 | 69.2% | 74.3% | | 52.6% |
| | FIT, 50-75, 1 | 16,105 | 839 | 969 | 1,829 | 10 | 35.1 | 12.3 | 391.1 | 285.6 | 56.3% | 66.5% | 92% | 47.4% |

AMR, adenoma miss rate; COL, colonoscopy; CRC, colorectal cancer; FIT, fecal immunochemical test; LY, life-years; LYG, life-years gained; N/A, not applicable.

**Table S6.** Screening outcomes in sensitivity analysis using updated input values. Outcomes are in scenarios of screening and follow-up/surveillance COL adenoma sensitivity. Adenoma sensitivity values by adenoma size and location and specificity by age for mt-sDNA and FIT were derived from a cross-sectional study (clinicaltrials.gov identifier, NCT01397747)[6]. Base-case adenoma sensitivity values for screening and follow-up/surveillance colonoscopy were derived from adenoma miss rate data in a meta-analysis published in 2019.[5] Results are per 1000 individuals screened with COL every 10 years, mt-sDNA every 3 years, or FIT every 1 year from ages 50–75 compared with no screening.

| Colonoscopy Performance Scenario | Screening Strategy | Stool Tests | Follow-up COLs | Surveillance COLs | Total COLs | Complications | CRC Cases | CRC Deaths | LY with CRC | LYG | Incidence Reduction | Mortality Reduction | % Threshold LYG Relative to COL | AMR, Weighted Mean |
|---|---|---|---|---|---|---|---|---|---|---|---|---|---|---|
| | No screening | 0 | 0 | 0 | 80 | 2 | 80.3 | 36.6 | 646.0 | 0.0 | 0.0% | 0.0% | | N/A |
| Scenario 1 | Colonoscopy 50-75, 10 | 0 | 0 | 1,658 | 4,131 | 15 | 15.2 | 5.9 | 169.1 | 344.6 | 81.1% | 84.0% | | 26.7% |
| | mt-sDNA, 50-75, 3 | 6,308 | 735 | 1,086 | 1,840 | 11 | 29.3 | 10.6 | 333.1 | 296.7 | 63.5% | 71.1% | 86% | 23.8% |
| | FIT, 50-75, 1 | 16,537 | 743 | 1,138 | 1,898 | 11 | 27.7 | 9.6 | 328.0 | 310.2 | 65.5% | 73.9% | 90% | 23.9% |
| Scenario 2 | Colonoscopy 50-75, 10 | 0 | 0 | 1,635 | 4,117 | 15 | 15.4 | 6.0 | 171.7 | 343.9 | 80.8% | 83.7% | | 29.3% |
| | mt-sDNA, 50-75, 3 | 6,298 | 735 | 1,101 | 1,854 | 11 | 28.9 | 10.4 | 328.4 | 298.6 | 64.1% | 71.7% | 87% | 21.8% |
| | FIT, 50-75, 1 | 16,516 | 742 | 1,153 | 1,911 | 11 | 27.1 | 9.4 | 322.3 | 311.8 | 66.2% | 74.3% | 91% | 21.8% |
| Scenario 3 | Colonoscopy 50-75, 10 | 0 | 0 | 1,607 | 4,097 | 15 | 15.8 | 6.1 | 175.0 | 342.3 | 80.4% | 83.4% | | 32.0% |
| | mt-sDNA, 50-75, 3 | 6,293 | 732 | 1,114 | 1,864 | 11 | 28.4 | 10.3 | 325.4 | 299.0 | 64.6% | 72.0% | 87% | 19.7% |
| | FIT, 50-75, 1 | 16,506 | 739 | 1,166 | 1,922 | 11 | 26.7 | 9.2 | 319.2 | 313.1 | 66.8% | 74.9% | 92% | 19.8% |
| Scenario 4 | Colonoscopy 50-75, 10 | 0 | 0 | 1,545 | 4,056 | 15 | 16.6 | 6.5 | 182.9 | 338.3 | 79.3% | 82.3% | | 37.7% |
| | mt-sDNA, 50-75, 3 | 6,284 | 729 | 1,136 | 1,882 | 11 | 27.7 | 10.0 | 320.3 | 302.2 | 65.5% | 72.7% | 89% | 16.4% |
| | FIT, 50-75, 1 | 16,479 | 738 | 1,191 | 1,945 | 11 | 25.9 | 9.0 | 313.5 | 315.0 | 67.7% | 75.5% | 93% | 16.4% |
| Scenario 5 | Colonoscopy 50-75, 10 | 0 | 0 | 1,520 | 4,040 | 15 | 17.4 | 6.8 | 192.0 | 334.0 | 78.4% | 81.5% | | 39.9% |
| | mt-sDNA, 50-75, 3 | 6,278 | 727 | 1,150 | 1,894 | 11 | 27.0 | 9.7 | 312.5 | 305.8 | 66.4% | 73.5% | 92% | 14.6% |
| | FIT, 50-75, 1 | 16,469 | 736 | 1,204 | 1,955 | 11 | 25.3 | 8.7 | 306.7 | 318.5 | 68.5% | 76.2% | 95% | 14.7% |
| Scenario 6 | Colonoscopy 50-75, 10 | 0 | 0 | 1,487 | 4,017 | 15 | 17.9 | 7.0 | 196.4 | 331.5 | 77.7% | 80.9% | | 42.7% |

| | | | | | | | | | | | | | | |
|---|---|---|---|---|---|---|---|---|---|---|---|---|---|---|
| | mt-sDNA, 50-75, 3 | 6,276 | 724 | 1,160 | 1,901 | 11 | 26.7 | 9.6 | 309.3 | 305.9 | 66.7% | 73.7% | 92% | 13.2% |
| | FIT, 50-75, 1 | 16,453 | 734 | 1,215 | 1,963 | 11 | 24.9 | 8.6 | 303.0 | 319.7 | 69.0% | 76.6% | 96% | 13.3% |
| Scenario 7 | Colonoscopy 50-75, 10 | 0 | 0 | 1,415 | 3,967 | 15 | 19.3 | 7.6 | 208.8 | 325.3 | 76.0% | 79.4% | | 48.4% |
| | mt-sDNA, 50-75, 3 | 6,266 | 724 | 1,176 | 1,916 | 11 | 26.1 | 9.4 | 306.7 | 308.0 | 67.4% | 74.4% | 95% | 10.7% |
| | FIT, 50-75, 1 | 16,442 | 732 | 1,231 | 1,977 | 11 | 24.5 | 8.4 | 300.0 | 320.5 | 69.5% | 76.9% | 99% | 10.8% |
| Scenario 8 | Colonoscopy 50-75, 10 | 0 | 0 | 1,376 | 3,941 | 15 | 20.7 | 8.2 | 226.0 | 318.0 | 74.2% | 77.6% | | 51.0% |
| | mt-sDNA, 50-75, 3 | 6,264 | 722 | 1,185 | 1,923 | 11 | 25.7 | 9.2 | 301.2 | 309.7 | 68.0% | 74.8% | 97% | 9.5% |
| | FIT, 50-75, 1 | 16,432 | 730 | 1,241 | 1,986 | 12 | 24.0 | 8.2 | 294.3 | 322.8 | 70.1% | 77.5% | 102% | 9.6% |
| Scenario 9 | Colonoscopy 50-75, 10 | 0 | 0 | 1,337 | 3,915 | 14 | 21.5 | 8.5 | 231.0 | 314.2 | 73.2% | 76.8% | | 53.8% |
| | mt-sDNA, 50-75, 3 | 6,260 | 722 | 1,191 | 1,929 | 11 | 25.6 | 9.2 | 300.6 | 310.9 | 68.1% | 75.0% | 99% | 8.5% |
| | FIT, 50-75, 1 | 16,429 | 729 | 1,248 | 1,991 | 11 | 23.8 | 8.2 | 293.8 | 322.6 | 70.3% | 77.6% | 103% | 8.6% |
| Scenario 10 | Colonoscopy 50-75, 10 | 0 | 0 | 1,255 | 3,858 | 14 | 23.4 | 9.3 | 247.6 | 306.4 | 70.9% | 74.7% | | 59.2% |
| | mt-sDNA, 50-75, 3 | 6,253 | 721 | 1,204 | 1,941 | 11 | 25.2 | 9.0 | 296.1 | 311.5 | 68.6% | 75.3% | 102% | 6.9% |
| | FIT, 50-75, 1 | 16,416 | 727 | 1,260 | 2,001 | 12 | 23.4 | 8.1 | 290.9 | 324.8 | 70.8% | 78.0% | 106% | 6.8% |

AMR, adenoma miss rate; COL, colonoscopy; CRC, colorectal cancer; FIT, fecal immunochemical test; LY, life-years; LYG, life-years gained; mt-sDNA, multitarget stool DNA test.

**Figure S1. Age-specific risks of complications from colonoscopy with polypectomy used in the analysis.** Complications include serious gastrointestinal events, other gastrointestinal events, and cardiovascular events. We assume that the only harms from screening arise from a colonoscopy with polypectomy, whether it be for screening, follow-up, or surveillance, or for the diagnosis of a symptomatic cancer. We assume no risk of harms from stool-based tests, nor from bowel preparation.[7] The risks of complications from colonoscopy are from an analysis by van Hees et al.,[8] which was an extension of an analysis by Warren et al.[9] In those studies, colonoscopy without polypectomy was not associated with an excess risk of complications, relative to a matched control group that did not have colonoscopy. Reproduced and adapted with permission from Knudsen et al, *JAMA.* 2016;315(23):2595-2609.[3]

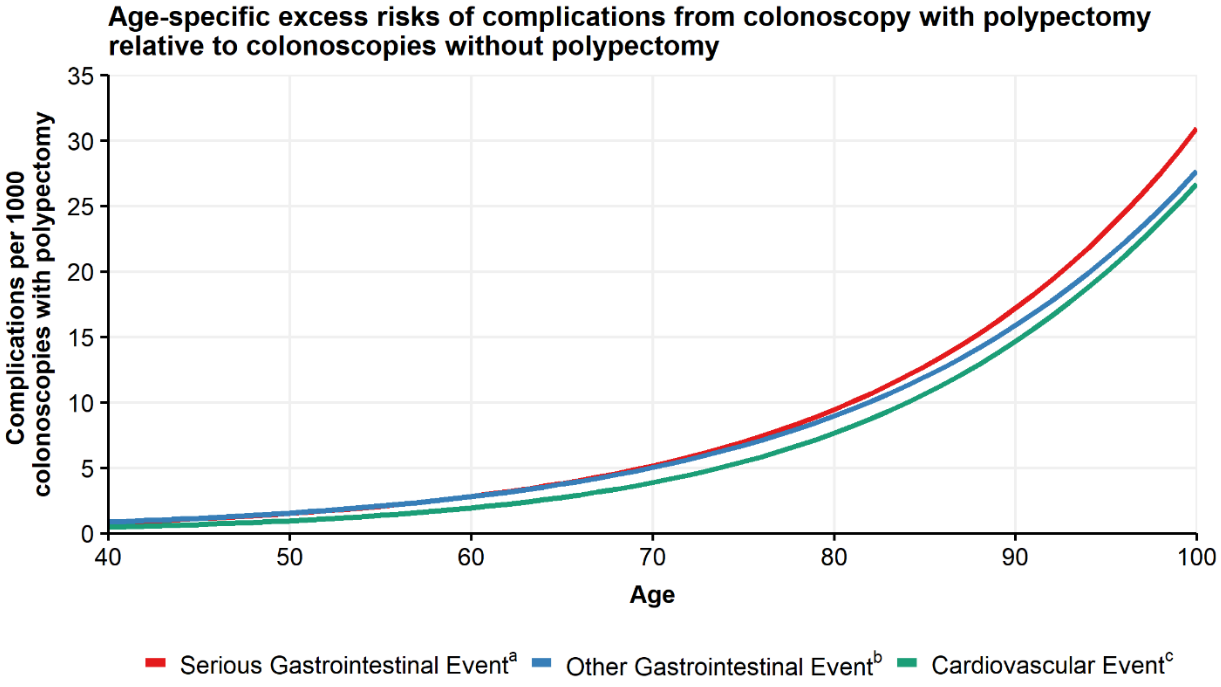


[a] Perforations, gastrointestinal bleeding or transfusions. Excess risk per colonoscopy with polypectomy = $1/[\exp(9.27953 - 0.06105 \times \text{Age}) + 1] - 1/[\exp(10.78719 - 0.06105 \times \text{Age}) + 1]$.

[b] Paralytic ileus, nausea and vomiting, dehydration, abdominal pain. Excess risk per colonoscopy with polypectomy $= 1/[\exp(8.81404 - 0.05903 \times \text{Age}) + 1] - 1/[\exp(9.61197 - 0.05903 \times \text{Age}) + 1]$.

[c] Myocardial infarction or angina, arrhythmias, congestive heart failure, cardiac or respiratory arrest, syncope, hypotension, or shock. Excess risk per colonoscopy with polypectomy $= 1/[\exp(9.09053 - 0.07056 \times \text{Age}) + 1] - 1/[\exp(9.38297 - 0.07056 \times \text{Age}) + 1]$.

**Figure S2.** Screening outcomes in sensitivity analysis using updated input values. A) Predicted life-years gained (LYG), B) reduction in CRC-related incidence and C) reduction in CRC-related mortality in scenarios of screening and follow-up/surveillance colonoscopy (COL) adenoma sensitivity. Adenoma sensitivity values by adenoma size and location and specificity by age for mt-sDNA and FIT were derived from a cross-sectional study (clinicaltrials.gov identifier, NCT01397747)[6]. Base-case adenoma detection sensitivity values for screening and follow-up/ surveillance colonoscopy were derived from adenoma miss rate data in a meta-analysis published in 2019.[5] Results are per 1000 individuals screened with COL every 10 years, mt-sDNA every 3 years, or FIT every 1 year from ages 50–75 compared with no screening.

A)

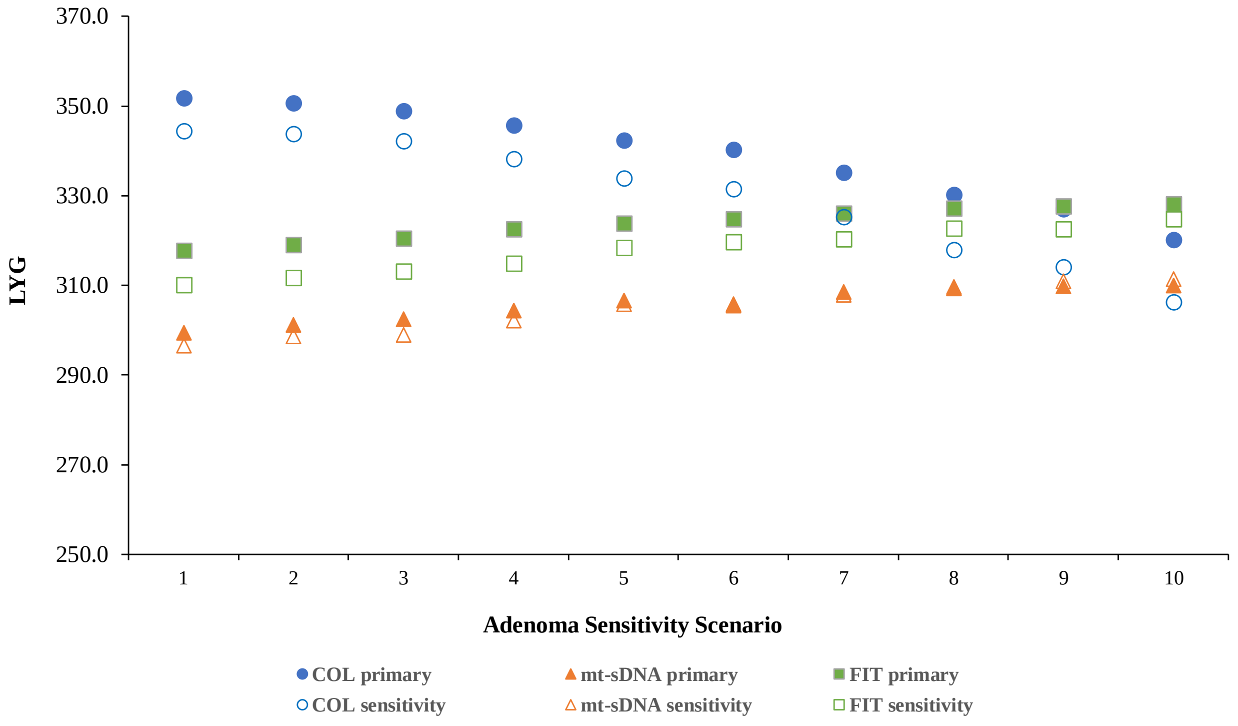


B)

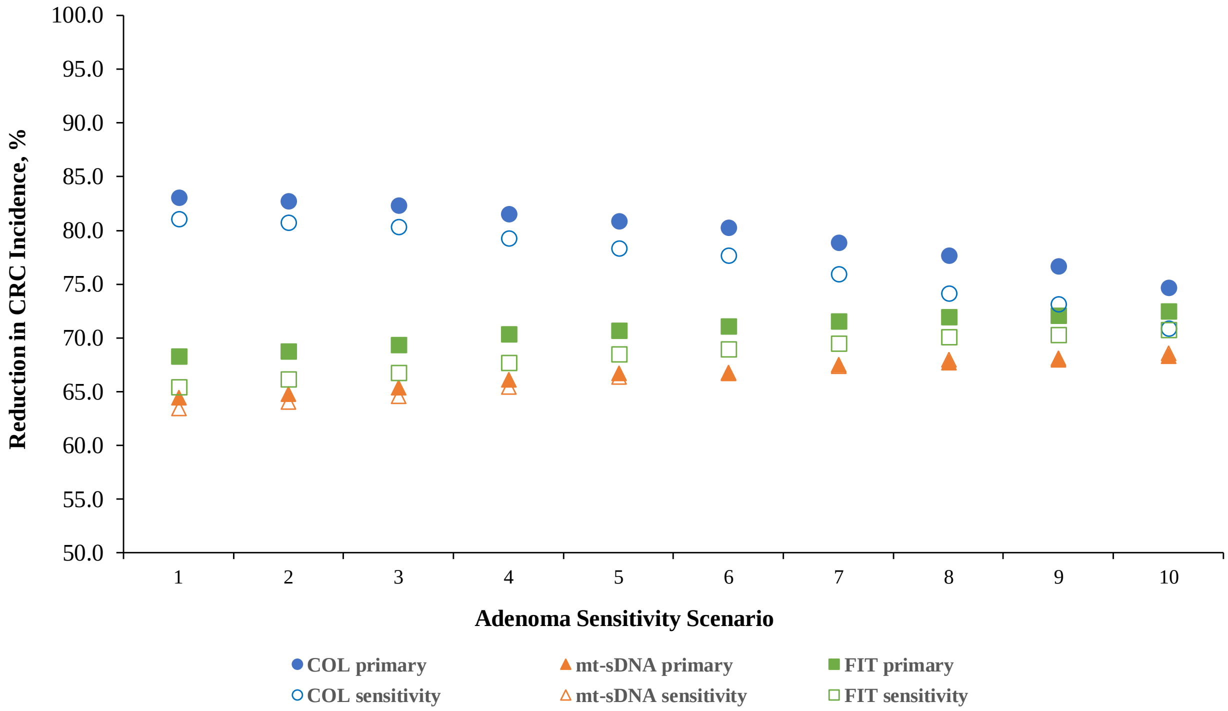

Reduction in CRC Incidence, %
100.0
95.0
90.0
85.0
80.0
75.0
70.0
65.0
60.0
55.0
50.0
1
2
3
4
5
6
7
8
9
10
Adenoma Sensitivity Scenario
COL primary
COL sensitivity
mt-sDNA primary
mt-sDNA sensitivity
FIT primary
FIT sensitivity


C)

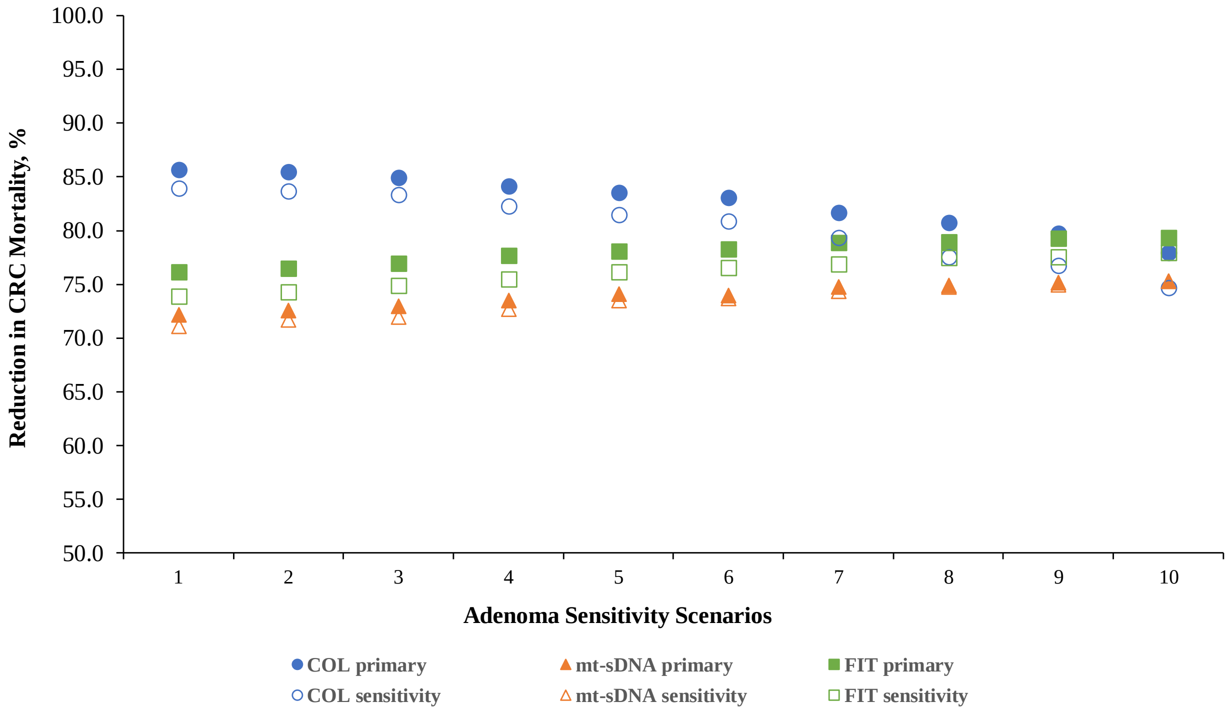

Reduction in CRC Mortality, %
100.0
95.0
90.0
85.0
80.0
75.0
70.0
65.0
60.0
55.0
50.0
1
2
3
4
5
6
7
8
9
10
Adenoma Sensitivity Scenarios
COL primary
COL sensitivity
mt-sDNA primary
mt-sDNA sensitivity
FIT primary
FIT sensitivity

**Figure S3.** Weighted mean adenoma miss rates (AMR) in sensitivity analysis using updated input values. Scenarios are of screening and follow-up/surveillance COL adenoma sensitivities for colonoscopy (COL) every 10 years, multitarget stool DNA test (mt-sDNA) every 3 years, or fecal immunochemical test (FIT) every year. Adenoma sensitivity values by adenoma size and location and specificity by age for mt-sDNA and FIT were derived from a cross-sectional study (clinicaltrials.gov identifier, NCT01397747)[6]. Base-case adenoma detection sensitivity values for screening and follow-up/ surveillance colonoscopy were derived from adenoma miss rate data in a meta-analysis published in 2019.[5]

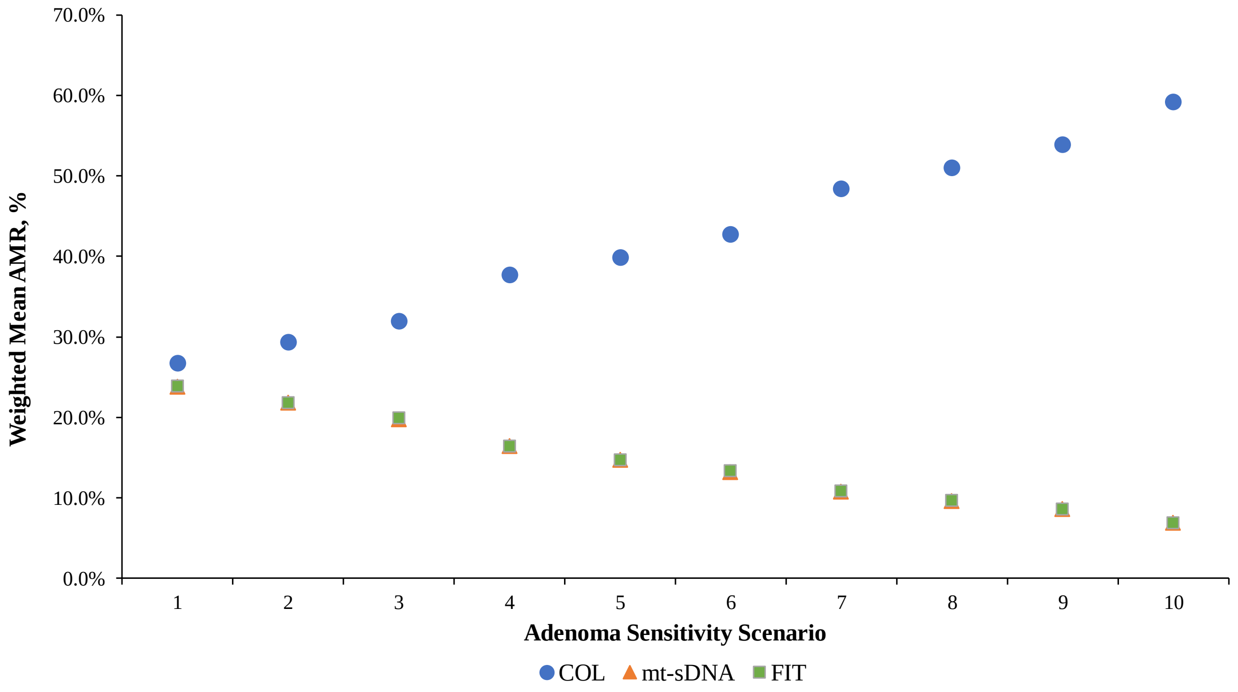
